# Identification of gut microbiome markers for schizophrenia delineates a potential role of *Streptococcus*

**DOI:** 10.1101/774265

**Authors:** Feng Zhu, Yanmei Ju, Wei Wang, Qi Wang, Ruijin Guo, Qingyan Ma, Qiang Sun, Yajuan Fan, Yuying Xie, Zai Yang, Zhuye Jie, Binbin Zhao, Liang Xiao, Lin Yang, Tao Zhang, Junqin Feng, Liyang Guo, Xiaoyan He, Yunchun Chen, Ce Chen, Chengge Gao, Xun Xu, Huanming Yang, Jian Wang, Yonghui Dang, Lise Madsen, Susanne Brix, Karsten Kristiansen, Huijue Jia, Xiancang Ma

## Abstract

Emerging evidence has linked the gut microbiota to schizophrenia. However, the functional changes in the gut microbiota and the biological role of individual bacterial species in schizophrenia have not been explored systematically. Here, we characterized the gut microbiota in schizophrenia using shotgun metagenomic sequencing of feces from a discovery cohort of 90 drug-free patients and 81 controls, as well as a validation cohort of 45 patients taking antipsychotics and 45 controls. We screened 83 schizophrenia-associated bacterial species and constructed a classifier comprising 26 microbial biomarkers that distinguished patients from controls with a 0.896 area under the receiver operating characteristics curve (AUC) in the discovery cohort and 0.765 AUC in the validation cohort. Our analysis of fecal metagenomes revealed that schizophrenia-associated gut–brain modules included short-chain fatty acids synthesis, tryptophan metabolism, and synthesis/degradation of neurotransmitters including glutamate, γ-aminobutyric acid, and nitric oxide. The schizophrenia-enriched gut bacterial species include several oral cavity-resident microbes, such as *Streptococcus vestibularis*. We transplanted *Streptococcus vestibularis* into the gut of the mice with antibiotic-induced microbiota depletion to explore its functional role. We observed that this microbe transiently inhabited the mouse gut and this was followed by hyperactivity and deficit in social behaviors, accompanied with altered neurotransmitter levels in peripheral tissues. In conclusion, our study identified 26 schizophrenia-associated bacterial species representing potential microbial targets for future treatment, as well as gut–brain modules, some of which may give rise to new microbial metabolites involved in the development of schizophrenia.

## Introduction

Schizophrenia is a severe psychiatric disorder associated with hallucinations, delusions, and thought disorders perturbing perception and social interaction^1^. The etiology of schizophrenia is not elucidated, but assumed to be multifactorial involving genetic and environmental factors. Abnormalities of neurotransmitter systems have been extensively studied especially focusing on aberration of signaling involving dopamine, glutamate, and γ-aminobutyric acid (GABA)^2–4^. Increasing evidence indicates that schizophrenia may be a systemic disorder with neuropsychiatric conditions in addition to psychosis^5^. Furthermore, the importance of inflammation^6^ and the involvement of the gastrointestinal system^7^ in schizophrenia have received attention.

The gut microbiota is reported to play an important role in neurogenerative processes, and perturbation of the microbiota and microbial products have been demonstrated to affect behavior^8–10^. Changes in the gut microbiota have been associated with neurological^11^ and neurodevelopmental disorders^12,13^, including schizophrenia^14^. It was recently reported that fecal transfer of the gut microbiota from patients with schizophrenia induces schizophrenia-associated behaviors in germ-free recipient mice accompanied with altered levels of glutamate, glutamine, and GABA in the hippocampus^13^. However, the identity and functionality of the specific bacteria responsible for mediating changes in the behavior of recipient mice are unknown^15,16^. Thus, the composition and functional capacity of the gut microbiota in relation to schizophrenia need to be systematically examined. Taxa and functional profiling of the microbiota composition is a premise for functional understanding of the gut microbiota^17^. Previous studies were based on 16S rRNA gene amplicon sequencing of relatively small cohorts^14,18–20^ and hence failed to provide detailed functional profiling of the gut microbiota in schizophrenia. Metagenomic shotgun sequencing combined with bioinformatics tools enables better characterization of the microbiota^21^, including a more accurate prediction of biological features of the microbes and their impacts on host physiology^22^.

Here, we carried out a metagenome-wide association study (MWAS) using 171 samples (90 cases and 81 controls) and validated the results with an additional 90 samples (45 cases and 45 controls). The functional changes within the schizophrenia gut microbiota were determined using pathway/module analysis based on the Kyoto Encyclopedia of Genes and Genomes (KEGG) and a recently developed gut–brain module (GBM) analysis of fecal metagenomes^23^. The functional roles of one particular schizophrenia-enriched gut bacterial species, *Streptococcus vestibularis*, were explored by transplanting this bacterium into the gut of the mice with antibiotic-induced microbiota depletion and observing its effects on animal behavior and brain neurochemicals.

## Results

### The gut microbiota differs between schizophrenic patients and healthy controls

We carried out shotgun sequencing on fecal samples from 90 drug-free patients and 81 healthy controls (for demographic and clinical characteristics see Supplementary Table 1-3) and obtained an average of over 11 gigabases (Gb) sequence data per sample and mapped the high-quality reads onto a comprehensive reference gene catalogue of 11.4 million genes^24^ (Supplementary Table 4).

The gut microbiota in schizophrenic patients showed greater α diversity at the genus level (*P* = 0.027, Wilcoxon rank-sum test), higher β diversity at the genus level (*P* < 0.001, Wilcoxon rank-sum test) and microbial gene level (*P* < 0.001, Wilcoxon rank-sum test) and comprised more genes compared with healthy controls (Supplementary Fig. 1). Out of a total of 360 metagenomic operational taxonomic units (mOTUs)^25^, 83 mOTUs showed significant differences in relative abundance between patients and controls (*P* < 0.05 and false discovery rate (FDR) = 0.136, Wilcoxon rank sum test and Storey’s FDR method; Supplementary Table 5a). After adjusting for BMI, age, sex, and diet, these 83 mOTUs were still significant (Supplementary Table 5a). The gut microbiota in schizophrenic patients harbored many facultative anaerobes such as *Lactobacillus fermentum, Enterococcus faecium*, *Alkaliphilus oremlandii*, and *Cronobacter sakazakii/turicensis*, which are rare in a healthy gut. Additionally, bacteria that are often present in the oral cavity, such as *Veillonella atypica*, *Veillonella dispar*, *Bifidobacterium dentium*, *Dialister invisus*, *Lactobacillus oris*, and *Streptococcus salivarius* were more abundant in patients with schizophrenia than in healthy controls, indicating a close association between the oral and the gut microbiota in schizophrenia.

**Figure 1.**
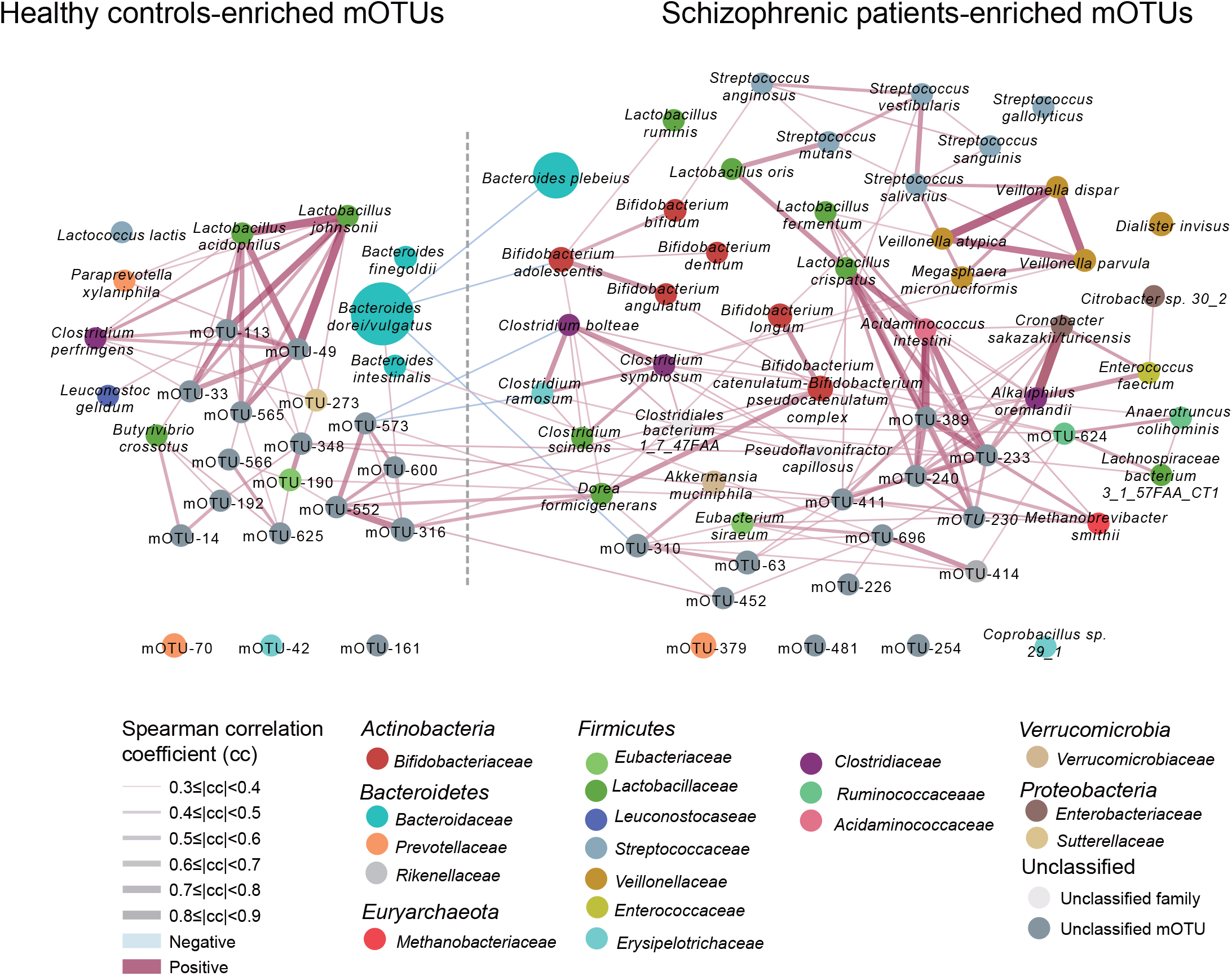
Network of mOTUs differentially enriched in healthy controls and schizophrenic patients. Node sizes reflect the mean abundance of mOTUs. mOTUs annotated to species are colored according to family (Red edges, Spearman’s rank correlation coefficient > 0.3, *P* < 0.05; blue edges, Spearman’s rank correlation coefficient < −0.3, *P* < 0.05;).

We then constructed a mOTU network to depict the co-occurrence correlation between the schizophrenia-associated gut bacteria (Fig. 1). Schizophrenia-enriched mOTUs (Wilcoxon rank-sum test) were more interconnected than control-enriched mOTUs (Spearman’s correlation coefficient < –0.3 or >0.3, *P* < 0.05). The mOTU species from the genera *Streptococcus* and *Veillonella* showed positive cross-correlations. Moreover, the majority of the species in these two clusters of correlated mOTUs originated from the oral cavity, suggesting that orally resident bacteria in a synergistic manner may colonize the gut in schizophrenic patients (Fig. 1 and Supplementary Table 5a).

Functional modules and pathways enriched in the gut microbiota of patients relative to controls were analyzed using the KEGG database (Supplementary Table 6). The relative enrichment of 579 KEGG modules and 323 KEGG pathways varied significantly between the two groups. Schizophrenia-depleted microbial functional modules included pectin degradation, lipopolysaccharide biosynthesis, autoinducer-2 (AI-2) transport system, glutamate/aspartate transport system, beta-carotene biosynthesis, whereas schizophrenia-enriched functional modules included methanogenesis, the gamma-aminobutyrate (GABA) shunt, and transport system of manganese, zinc, and iron (Supplementary Table 6).

### Neuroactive potential of schizophrenia-related bacterial species

We next compared the altered microbial neuroactive potential of the gut microbiota of schizophrenic patients with the controls at the species level using the method reported by Valles-Colomer et al.^23^. We mapped the metagenomic data of the 171 samples to a genome database including the 42 microbial species that were detected based on the 83 schizophrenia-associated mOTUs using PanPhlAn^26^ and calculated the prevalence of species-level microbes. We then determined whether the abundance of 56 previously reported gut-brain modules (GBMs)^23^, present in each microbial species, varied significantly between schizophrenic patients and controls. The GBM set of each microbial species was obtained by cross-checking GBM-related genes and the species gene repertoires (Supplementary Table 7a). The frequency of the occurrence of each GBM within each species was compared between patients and controls using a Chi-squared test (Supplementary Table 7b). Schizophrenia-associated GBMs included short-chain fatty acid synthesis (acetate, propionate, butyrate, and isovaleric acid), tryptophan metabolism, and the synthesis of several neurotransmitters, such as glutamate, GABA, and nitric oxide (Fig. 2).

**Figure 2.**
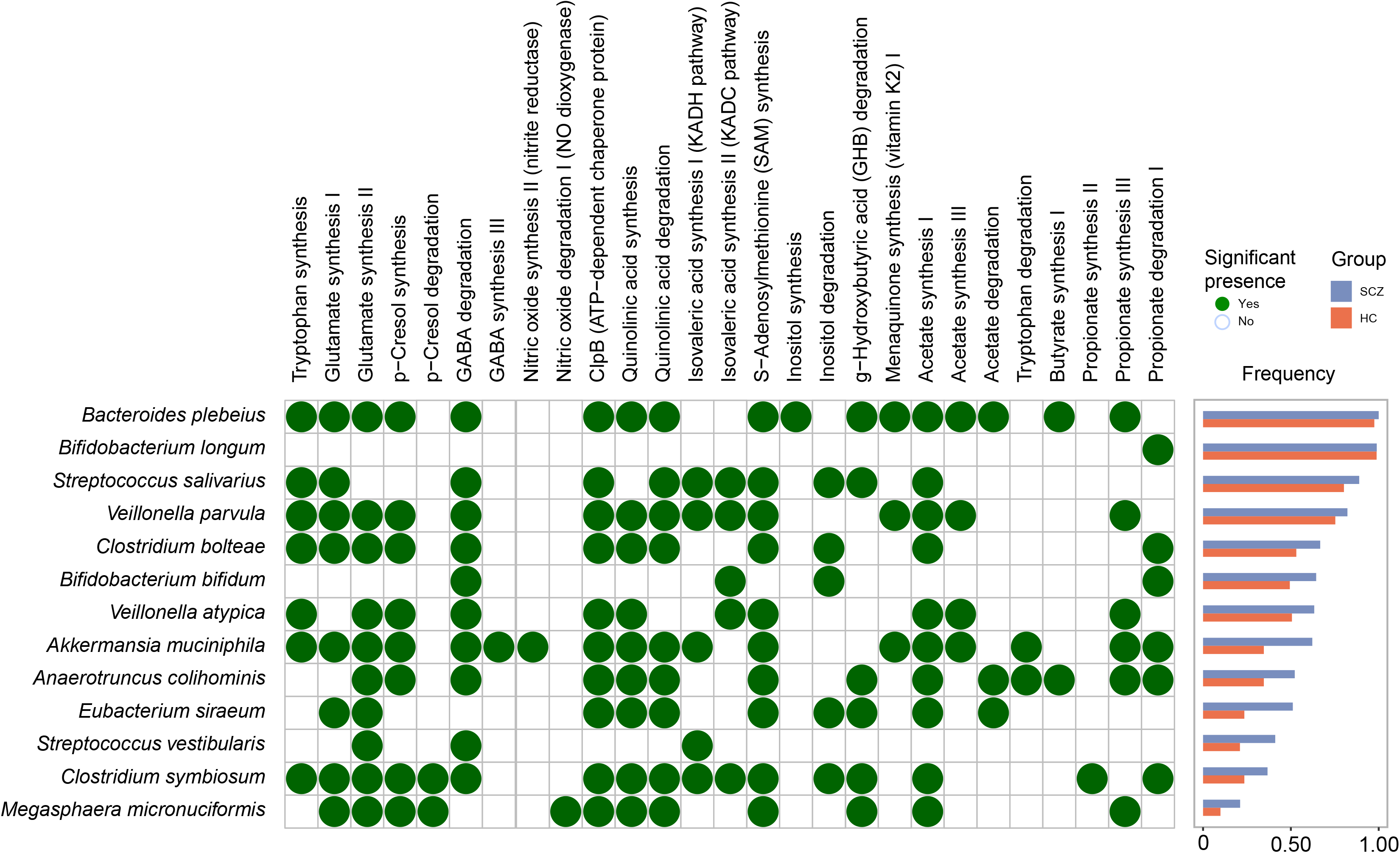
The gut-brain modules present in schizophrenia-associated bacterial species. A green dot indicates a statistically significant association between a gut-brain modules present in schizophrenia-associated bacterial species and a metabolite. No dot represents a non-significant association or a non-existent association. The difference in relation to presence between schizophrenic patients and heathy controls was calculated (Chi-square test, *P* < 0.05). The bar plot shows the frequency of each bacterial species present in schizophrenic patients (SCZ, blue bar) and healthy controls (HC red bar), respectively.

We chose to validate the presence of the GBM associated with tryptophan metabolism in schizophrenia, as tryptophan metabolism also is modulated by the gut microbiota and implicated in schizophrenia pathogenesis^27,28^. Hence, serum tryptophan metabolites were measured in patients and controls and correlated with the presence of tryptophan modules in the gut microbiota. In agreement with the higher abundance of tryptophan metabolisms related GBMs, we observed lower serum tryptophan levels and higher kynurenic acid (KYNA) levels in schizophrenic patients(Supplementary Fig. 2a, c). Moreover, serum tryptophan levels were negatively correlated with the abundances of 38 bacterial species enriched in schizophrenic patients and positively correlated with six bacterial species enriched in controls (Supplementary Fig. 2d). Similarly, serum KYNA levels were positively correlated with 10 schizophrenia-enriched bacterial species and negatively correlated with 3 control-enriched bacterial species (Supplementary Fig. 2d). Thus, an altered gut microbiota may be associated with changes in serum levels of tryptophan and KYNA in schizophrenia.

### Microbial species-based biomarkers for schizophrenia

To identify novel gut bacterial biomarkers and evaluate their diagnostic values for schizophrenia, we first constructed a set of random forest disease classifiers based on gut mOTUs. We performed a five-fold cross-validation procedure ten times on 90 patients and 81 controls. Twenty six gut mOTUs reached the lowest classifier error in the random forest cross validation, and the ROC AUC score of the model was 0.896 (Fig 3a, b). This microbial based classifier was not significantly influenced by age, gender, BMI, and diet style (Supplementary Table 8). This discriminatory model was validated on an additional validation cohort consisting 45 patients taking antipsychotics and 45 controls (Supplementary Table 9). The model still distinguished patients from controls with an ROC AUC score of 0.765. Among the 26 microbial biomarkers, 11 bacterial species with taxonomic identity were significantly enriched in schizophrenia, namely *Akkermansia muciniphila*, *Bacteroides plebeius*, *Veillonella parvula*, *Clostridium symbiosum*, *Eubacterium siraeum*, *Cronobacter sakazakii/turicensis*, *Streptococcus vestibularis, Alkaliphilus oremlandii*, *Enterococcus faecium*, *Bifidobacterium longum*, and *Bifidobacterium adolescentis*. Some of these microbial biomarkers were significantly associated with symptom severity, cognitive performance, and diagnosis (Fig 3c). We next performed metagenomic analysis on the fecal samples from 38 of the 90 patients after 3-months of treatment (27 with risperidone and 11 with other antipsychotics, shown in Supplementary Table 1). The psychotic symptoms and cognitive impairment improved greatly along with treatment (Supplementary Fig. 3). However, only approximately half of the microbial biomarkers returned to the levels in controls after treatment (Fig. 3d). As the sample size of the follow-up patients was smaller, the statistical significance threshold was increased from 0.05 to 0.1. Of the 26 identified microbial biomarkers, 20 biomarkers remained significantly changed between 81 controls and 38 baseline patients (*P* < 0.1, FDR = 0.44, Benjamini and Hochberg method, Fig. 3d). After 3-months of treatment, the abundances of 12 of these 26 biomarkers remained significantly changed compared with 81 controls (*P* < 0.1, FDR = 0.33, Benjamini and Hochberg method, Supplementary Table 10). Pair-wise comparison of all gut mOTUs for the treatment effect in the follow-up patients revealed 48 differentially abundant bacterial species (*P* < 0.05 and FDR = 0.420, Paired Wilcoxon rank sum test; Benjamini and Hochberg method, Supplementary Table 11). However, only 5 of the 48 differentially abundant species were included in the 26 microbial biomarkers used for diagnosis. This result suggests that antipsychotic treatment influences the gut microbiota, but does not completely restore the altered microbiota associated with schizophrenia.

**Figure 3.**
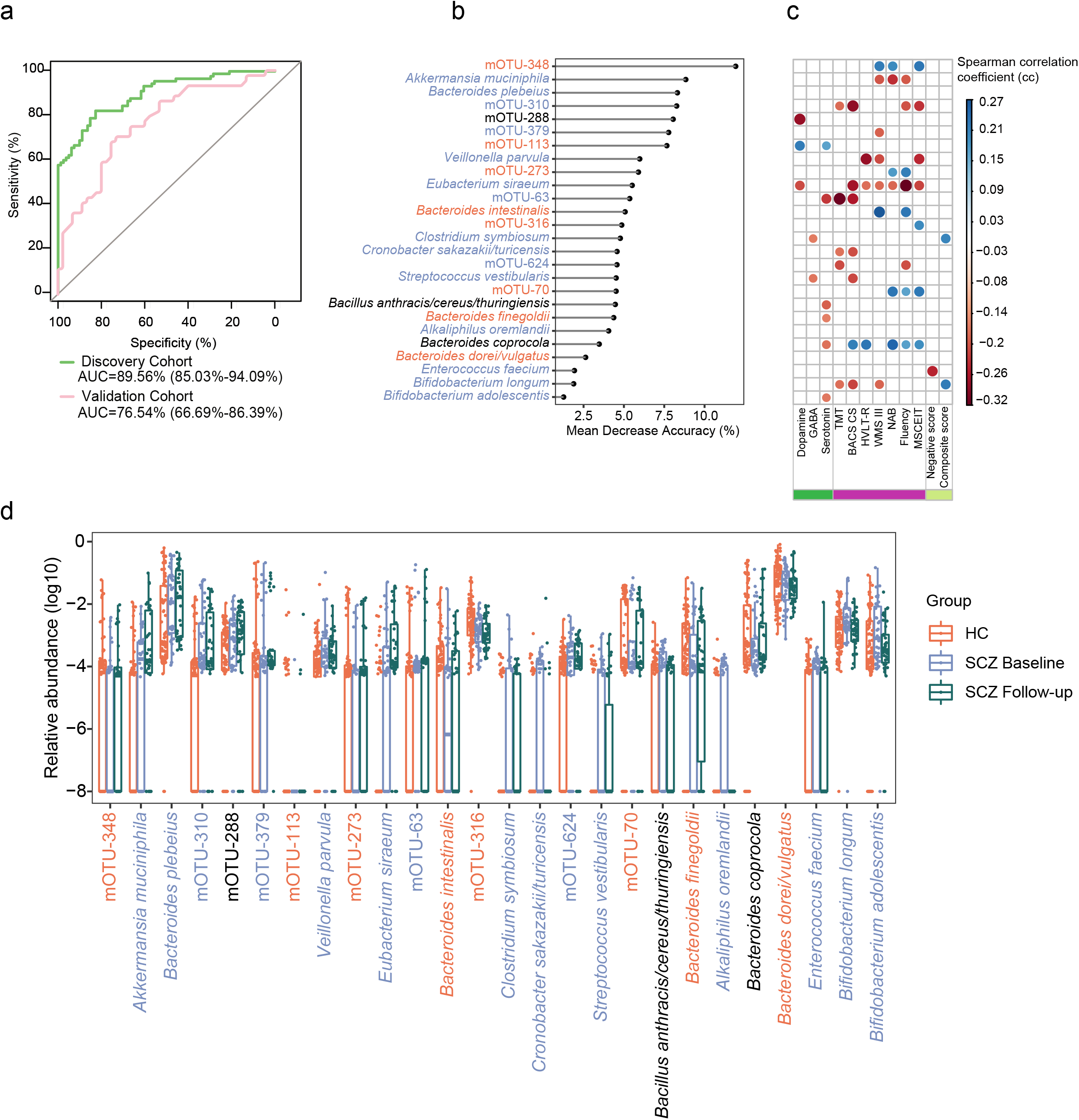
Gut microbiome-based dicrimination between schizophrenic patients and healthy controls. (a) Receiver operating curves (ROC) according to 171 samples of the discovery set (green line) and 90 independent validation samples (pink line) calculated by cross-validated random forest models. Area under ROC (AUCs) and the 95% confidence intervals (in parentheses) are also shown. (b) The 26 mOTUs with most weight to discriminate schizophrenic (SCZ) patients and healthy controls (HC) were selected by the cross-validated random forest models. The length of line indicates the contribution of the biomarkers to the discriminative model. The color of each mOTU indicates its enrichment in schizophrenic patients (blue) or healthy controls (purple) or no significant direction (black), respectively. (c) Spearman’s correlation of 26 mOTUs classifiers with three types of neurotransmitter in serum (green), seven types of cognitive function evaluated using the MATRICS Consensus Cognitive Battery (purple), and with the positive score and the negative score of the positive and negative syndrome scale (blue). Only significant associations are displayed with correlation coefficient (*P* value < 0.05). (d). The relative abundance (log_10_) of 26 mOTU classifiers in 90 HCs and 38 SCZ patients at baseline and on a follow-up (3 months later). GABA: 4-aminobutyric acid; TMT: Trail Making Test; BACS SC: Brief Assessment of Cognition in Schizophrenia; Fluency: Category Fluency in Animal Naming; WMS-III: Wechsler Memory Scale-Third Edition for working memory; HVLT-R: Hopkins Verbal Learning Test-Revised for visual learning; NAB: Neuropsychological Assessment Battery for reasoning and problem solving; MSCEIT: Mayer-Salovey-Caruso Emotional Intelligence Test for social cognition.

### Streptococcus vestibularis induced schizophrenia-like behaviors in mice

*Streptococcus* (*S.*) *vestibularis* was identified as a diagnostic marker associated with serum GABA, tryptophan, KYNA, and the Brief Assessment of Cognition in Schizophrenia (BACS) scores in MATRICS Consensus Cognitive Battery (MCCB) test (Fig. 3 and Supplementary Fig. 2). Moreover, *S. vestibularis*, present in the gut of a number of schizophrenic patients, was predicted to have GBMs related to glutamate synthesis, GABA degradation, and Isovaleric acid synthesis I (Fig. 2). As some pathogenic species of *Streptococcus* are known to enter the brain^29^, and have been implicated in pediatric acute-onset neuropsychiatric syndrome^30,31^, we asked if *S. vestibularis* might play a role in the pathophysiology of schizophrenia. Hence, we transplanted *S. vestibularis* ATCC 49124, using oral gavage and drinking water, into C57BL/6 mice after antibiotics-based microbiota depletion (Supplementary Fig. 4). Another strain of streptococcus, *S. thermophilus* ST12, which is widely present in the human gut, was used as a bacterial control. Behavioral tests were performed to evaluate the effect of *S. vestibularis* transplantation (Fig. 4a). Quantitative polymerase chain reaction (q-PCR) used to quantify the 16S rRNA gene of *S. vestibularis* and *S. thermophilus*, revealed that their concentration increased by 4,164- and 6,183-fold immediately after transplantation and remained at a 31.2- and 58.1-fold increase after the behavioral tests as compared to the control mice (Supplementary Fig. 5). Compared to control mice gavaged with buffer or with S. thermophilus, the S. vestibularis-treated mice had increased their total distance traveled and times of rearing during a 30-minute open-field test (Fig. 4b in this figure, it is better to put S. thermophilus next to control since S. thermophilus is also a control, d). They continued their hyperlocomotion after the 10-minute habituation period and showed no obvious decline in locomotion activity after a period of 30-minutes (Fig. 4c). In the three-chamber social test, the S. vestibularis mice displayed obvious deficits in sociability and social novelty, as they were much less sociable and avoided social novelty (Fig. 4e-g). However, in Barnes maze, elevated plus maze, and tail suspension test, the mice transplanted with *S. vestibularis* displayed spatial memory function, depressive state, and anxiety levels similar to either saline or *S. thermophilus*-treated mice (Supplementary Fig. 6). There were no significant changes in body weight, systemic pro-inflammatory cytokines, and endotoxin, and HPA axis hormones among the S. vestibularis-treated, S. thermophilus-treated, and control mice (Supplementary Fig. 7 a-i).

**Figure 4.**
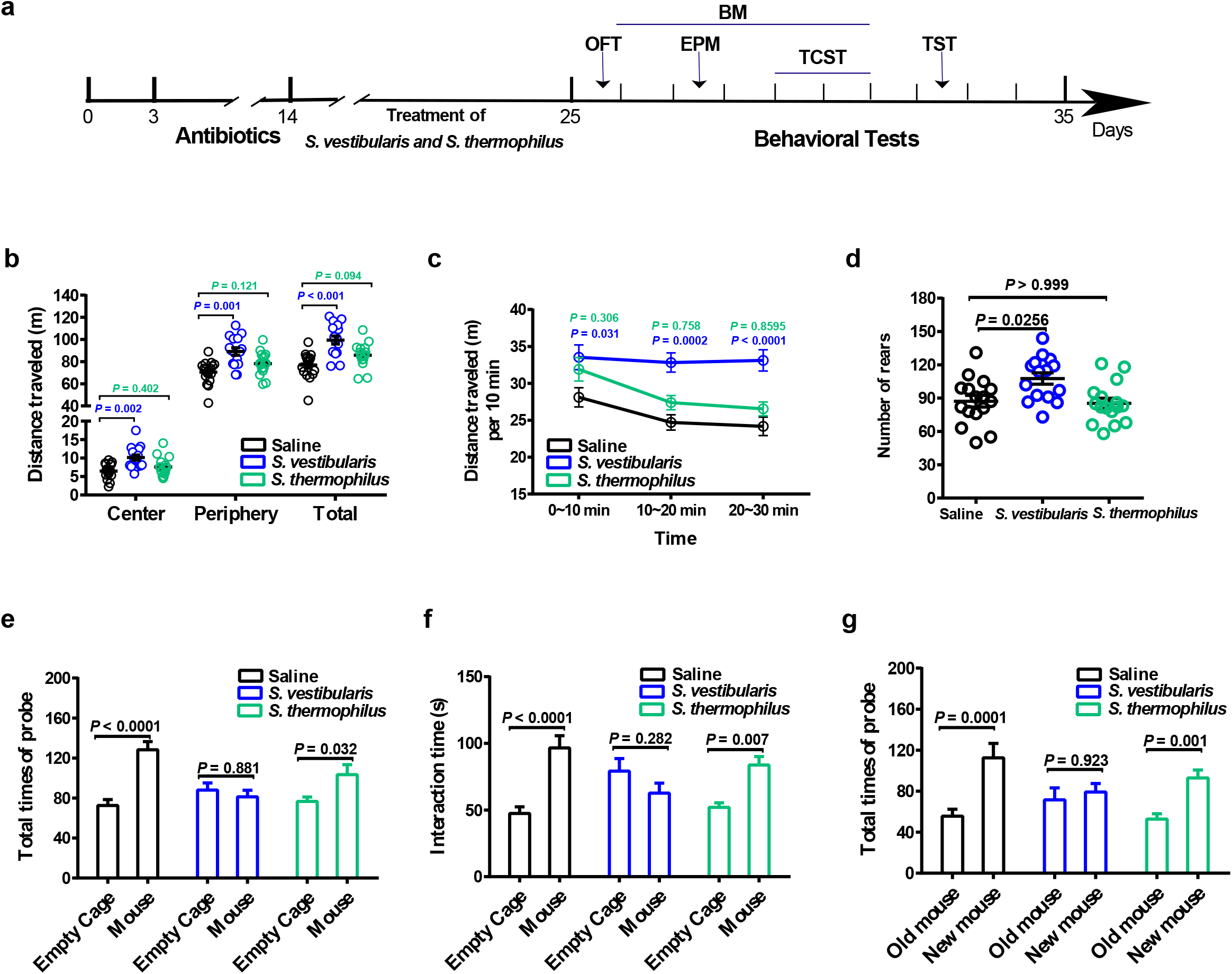
*Streptococcus vestibularis* induces hyperkinetic behavior and impaired social interaction in mice. (a) Schematic diagram of bacterial transplantation and behavioral test. The gut microbiota of mice was depleted by antibiotics, and then *Streptococcus vestibularis*, *Streptococcus thermophilus*, or their storage buffer (saline) was administered via oral gavage and drinking water. A series of behavioral tests were carried out after administration of bacteria. The data are representative of two independent experiments. Open field test (OFT), three-chamber social test (TCST), elevated plus maze (EPM), Barnes maze (BM), and the tail suspension test (TST) (for a detailed description, please consult Supplementary Figure 6). (b) The cumulative distance (meters) in different zones in 30-minute OFT in the three groups of mice (in periphery: F_(2,46)_ =11.12, *P* = 0.0001; in center: F_(2,46)_ = 9.221, *P* = 0.0004; total distance: F_(2,46)_=13.69, *P* < 0.0001). (c) the cumulative distance (meters) in every 10-minute time interval of OFT traveled by *Streptococcus vestibularis*-treated mice compared to that traveled by *Streptococcus thermophilus*-treated mice and control mice. Time effect: F_(2,92)_ = 9.816, *P* = 0.0001; group effect: F_(2,46)_ = 13.69, *P* < 0.0001; time × group effect: F_(4,92)_ = 1.808, *P* = 0.134. (d) The number of rearing in *Streptococcus vestibularis*-treated mice compared with *Streptococcus thermophilus*-treated mice and saline-treated mice during a 30-minute OFT (*P* = 0.017). (e-g) TCST comparing sociability of *Streptococcus vestibularis*-treated mice to that of *Streptococcus thermophilus*-treated mice and control mice. The results show that *Streptococcus thermophilus*-treated mice and saline-treated mice display obvious sociability, i.e. demonstrate an increase in the number of times of probing a mouse (e, *P* < 0.0001) and spend longer time interacting with a mouse (f, *P* = 0.002) compared to an empty cage, and obvious social novelty, i.e. spend longer time interacting with an unacquainted mouse (new mouse; g, *P* = 0.005, DF = 90, t = 4.277) in comparison with an acquainted mouse. However, these types of social behaviors were not observed in *Streptococcus vestibularis*-treated mice (for sociability: e, *P* = 0.881, DF = 90, t = 0.665; f, *P* = 0.282, DF = 90, t = 1.639; for social novelty, g, *P* = 0.923; DF = 90, t = 0.564). Data are presented as Means ± SEM (n = 15~6 per group). The circle represents one value from individual mice (b and c). *P* values were determined one-way analysis of variance (ANOVA) (b), repeated measure two-way ANOVA followed by Sidak’s multiple comparisons test (c), Kruskal-Wallis test followed by Dunn’s multiple comparisons test (d), or two-way (ANOVA) followed by Sidak’s multiple comparisons test (e to g). Blue *P* value indicates the comparison between the *Streptococcus vestibularis*-treated and the saline-treated mice and green *P* value indicates the comparison between *Streptococcus thermophilus*-treated and saline-treated mice (b, c).

We then compared the transcriptome and neurotransmitter levels between *S. vestibularis*-treated and saline-treated mice in their peripheral tissues and brain. *S. vestibularis* treated mice had significantly lower levels of dopamine in serum, intestinal contents, and colonic tissue, as well as decreased GABA levels in the intestinal contents immediately after the transplantation, but these effects disappeared after 10 days of post-transplantation (Supplementary Fig. 8b, e, h, and f). Intestinal contents of S. vestibularis-treated mice showed increased levels of 5-HT throughout the behavioral tests (Supplementary Fig. 8d). S. vestibularis transplantation did not induce obvious inflammatory cell infiltration (Supplementary Fig. 6j) but induced changes in the expression of numerous immune/inflammation-related genes in the intestine when compared with saline gavage (Supplementary Table 12 a-b). By gene enrichment anslysis, we found these genes were enriched in cytokine-cytokine receptor interaction, chemokine signaling pathway, leukocyte trans-endothelial migration, complement and coagulation cascades, antigen processing and presentation, intestinal immune network for IgA production, and inflammatory bowel disease (Supplementary Fig. 9a-b). These results suggested that *S. vestibularis* may influence gut immune homeostasis. In the brain, the levels of neurotransmitters were not affected by transplantation with *S.* vestibularis and only tryptophan decreased in the prefrontal cortex of S. vestibularis-treated mice (Supplementary Table 13). However there are 354, 540, and 470 significantly differentially expressed gene in the PFC, striatum, and hippocampus between *S. vestibularis*-treated and saline treated mice (Supplementary Table 12 c-d). The pathways influenced by these differentially expressed genes included defense responses and immune-regulating pathways like in the gut, as well as peroxisome proliferator-activated receptor signaling pathway, steroid biosynthesis, tyrosine, and tryptophan metabolism (Supplementary Fig. 9c-e).

## Discussion

We here used a combination of MWAS and a functional-based approach to systematically screen for schizophrenia-associated gut microbes and identified a number of schizophrenia-associated GBMs, expanding our insight into functional changes characterizing the gut microbiota of schizophrenic patients. Consistent with the MWAS results and the inferred microbial functions, the schizophrenia-associated bacterium *Streptococcus vestibularis* was shown in mice to have a functional neuroactive potential^23^ associated with changes in animal behaviors.

Poor reproducibility is common in microbiome research^32^. In the present study, a diagnostic model of 26 microbial biomarkers obtained using a discovery cohort, which included drug-free patients, was well validated in a testing cohort, which included patients taking antipsychotics. We chose to investigate drug-free patients in the discovery set in order to eliminate the possible effects of antipsychotics on the gut microbiota and identify gut bacteria possibly involved in the development of schizophrenia. The validation of the initial findings in patients taking antipsychotics demonstrated that these microbial markers are, to a certain extent, independent of antipsychotics. Follow-up analysis also revealed that 20 of the 26 identified microbial biomarkers in the diagnostic model remained the same over a treatment duration of three months. Therefore, most microbial biomarkers for schizophrenia seem stable and are not sensitive to current antipsychotics.

Analysis of the bacterial V3-V4 region of 16S rRNA regions has a limited resolution in terms of identification of bacterial species^17,25,33^. Current 16S rRNA gene amplicon sequencing generally capture reliable taxonomic classification at the genus level^33^. However, several recent analyses indicate that many taxonomic associations might be presented only at levels subordinate to species^17,34,35^. Accordingly, most of the schizophrenia-associated microbial species revealed by the MWAS results were not identified in previous studies by using 16S rRNA gene sequencing^14,18,36^. There are more overlaps of findings at genus level between our findings and the previous studies (Supplementary Table 5b), including 6 in the study of Zheng et al. (*Acidaminococcus, Akkermansia, Alistipes, Citrobacter, Dialister, Veillonella*)^14^, 1 in the study of Schwarz et al. (*Lactobacillus*)^36^, 1 in the study of Shen et al. (*Methanobrevibacter*)^18^. Surprisingly, gut microbiota diversities based on genus level taxonomy and annotated genes were higher in schizophrenic patients compared to controls. In accordance with our data, both α diversity and β diversity showed an increase in the blood microbiota of schizophrenic patients^37^. The microbes in blood are thought to originate from the gut as well as from the oral cavities^38,39^. Moreover, the increased diversity of the blood microbiota may be due to the nonspecific overall increased microbial burden in schizophrenia^37^, which is supported by our observed increased microbial gene number in patients’ gut. The considerable heterogeneity of the etiology and clinical manifestation of schizophrenia^40,41^ may be implicated in such an increase in the microbiota diversity. Another notable feature of the gut microbiota in schizophrenia is the significant enrichment of oral cavity resident bacteria. Increased bacterial translocation due to a leaky gut and innate immune imbalance are both presented in patients with schizophrenia^42,43^. Furthermore, gastrointestinal inflammation due to a dysfunctional immune response to pathogen infection and food antigens is also prevalent in schizophrenia^44,45^. These intestinal pathological conditions may disrupt the mucosal barrier and decrease immune surveillance towards foreign microbes^7^, increasing the possibility of observing oral bacteria in the gut.

The composition of human gut microbiota is linked to schizophrenia^14,18,36^, but knowledge of individual microbial species is needed to decipher their biological role^46^. We still do not completely understand the functions of most of the schizophrenia-associated microbes identified here in the gut or their biological roles in schizophrenia. Intriguingly, some schizophrenia-enriched bacterial species in the present study are also over-presented in subjects with metabolic disorders and atherosclerotic cardiovascular diseases^47–50^. Schizophrenic patients are more likely to develop obesity, hyperglycemia/diabetes, hypertension, and cardiovascular disease. Additionally, some of these risks are independent of the effects of antipsychotic administration and healthy lifestyle choices^51–54^. Several prenatal and early-life risk factors are shared by schizophrenia, metabolic disorders, and cardiovascular diseases such as prenatal famine, postnatal growth restriction, the quality of fetal growth, and low birth weight^55–61^. Gut microbes enriched in both schizophrenia and metabolic disorders/cardiovascular disease may account for the increased risk of these comorbidities in schizophrenia. Moreover, new evidence also indicates that metabolic disorders in schizophrenia are not only comorbidities, but also affect pathogenesis, such as the manifestation of negative symptoms^62^, cognitive function^63^, and brain white matter disruption^64^. Treating metabolic disorders via physical activity and psychosocial and dietary interventions, is also an effective approach to improve the symptoms of schizophrenia^63^. Therefore, manipulation of gut microbes may have double therapeutic potential for both metabolic disorders and schizophrenia.

Identifying the mediators of gut–brain microbiota communication affected by specific microbial metabolites is a key step to elucidate the mechanism behind the regulation of the host’s behaviors and psychology by gut microbial species. A recently published database of manually curated GBMs provides a powerful tool for functional interpretation of metagenomes in a microbiota–gut–brain context^23^. Using this database, we identified 27 schizophrenia-associated GBMs, which give clues about how the gut microbiota might modulate the pathophysiology of schizophrenia. Among these GBMs, a few well-known molecular entities associated with schizophrenia were covered, such as several kinds of neurotransmitters^2–4^ and tryptophan metabolites^65,66^. Furthermore, some microbes presenting these GBMs were significantly associated with the serum levels of several neurotransmitters and tryptophan metabolites. Parallel to the present study, our previous animal study demonstrated that transplantation of fecal microbiota from drug-free patients with schizophrenia into specific pathogen-free mice could cause schizophrenia-like behavioral abnormalities and dysregulated kynurenine metabolism^67^. The consistent findings of altered tryptophan-kynurenine metabolism revealed by human serum metabolite analysis, microbiota-based GBM prediction, and mouse studies suggest that this pathway is an important link between schizophrenia and gut microbiota dysbiosis. Of note, transplantation of one bacteria, *S. vestibularis ATCC 49124*, possessing a GBM involved in GABA degradation induced abnormal behaviors in the recipient mice. Importantly, the level of GABA in intestinal content of *S. vestibularis*-treated mice significantly decreased compared with saline-treated mice, which is in accordance with the inferred GBM and GABA degradation. The schizophrenia-enriched *S. vestibularis* contributed to the expression of two types of schizophrenia-relevant behaviors (hyperactivity and impaired social behaviors) in mice. To the best of our knowledge, this is the first study which aims to determine the functional roles of a single bacterium associated with schizophrenia in disease pathogenesis. Although the biological mechanisms underlying the effects of S. vestibularis are still unclear, our data indicate profound influences of this microbe on brain neurotransmitters. In conclusion, our study identified a number of schizophrenia-associated bacterial species representing potential microbial targets for future treatment and GBMs of which some may support the synthesis of new microbial metabolites of importance for the development of schizophrenia

## Acknowledgements

This study was supported by the Clinical Research Award of the First Affiliated Hospital of Xi’an Jiaotong University (No. XJTU1AF-CRF-2016-005), Shenzhen Municipal Government of China (No. JCYJ20170817145523036 and DRC-SZ [2015]162), Innovation Team Project of Natural Science Fund of Shanxi Province (2017KCT-20), and Key Program of Natural Science Fund of Shanxi Province (2018ZDXD-SF-036).

## Author contribution

X. M., H. J., R. G., F. Z., and Z. J. conceived the study. F. Z., Z. Y., F. Z., L. Y, B. Z., and Q.M. performed mice experiments. W. W., Q. M., Y. F., L. G., Y. D., Y. C., C. C., C. G., and X.M. recruited volunteers and collected samples for the study. W. W., Q. M., Y. F., Z. Y., L. Y., and F. Z. collected the human fecal and blood samples. Z. Y., L. Y, and F. Z. analyzed the human serum. R. G.,Y. J., Q. W., Q. S., Y. X, and B. L. performed bioinformatics analyses. K. K., S. B., L. M, and B. L. advised on the mice experiments. F. Z., R. G.,H. J., Y. J., and Q. W. interpreted the results and wrote the manuscript with extensive revision performed by L.M., S.B.,K.K and Y.X. All authors contributed to the final revision of the manuscript.

## Data availability

Metagenomic sequencing data for all samples have been deposited in the European Nucleotide Archive (ENA) database under accession identification code ERP111403.

